# Loss of CG methylation in *Marchantia Polymorpha* causes disorganization of cell division and reveals unique DNA methylation regulatory mechanisms of non-CG methylation

**DOI:** 10.1101/363937

**Authors:** Yoko Ikeda, Ryuichi Nishihama, Shohei Yamaoka, Mario A. Arteaga-Vazquez, Adolfo Aguilar-Cruz, Daniel Grimanelli, Romain Pogorelcnik, Robert A. Martienssen, Katsuyuki T. Yamato, Takayuki Kohchi, Takashi Hirayama, Olivier Mathieu

## Abstract

DNA methylation is an epigenetic mark that ensures silencing of transposable elements (TEs) and affects gene expression in many organisms. The function of different DNA methylation regulatory pathways has been largely characterized in the model plant *Arabidopsis thaliana*. However, far less is known about DNA methylation regulation and functions in basal land plants. Here we focus on the liverwort *Marchantia polymorpha*, an emerging model species that represents a basal lineage of land plants. We identified Mp*MET*, the *M. polymorpha* orthologue of the *METHYLTRANSFERASE 1* (*MET1*) gene required for maintenance of methylation at CG sites in angiosperms. We generated Mp*met* mutants using the CRISPR/Cas9 system, which showed a significant loss of CG methylation and severe morphological changes and developmental defects. The mutants developed many adventitious shoot-like structures, suggesting that Mp*MET* is required for maintaining differentiated cellular identities in the gametophyte. Numerous TEs were up-regulated, even though non-CG methylation was highly increased at TEs in the Mp*met* mutants. Closer inspection of CHG methylation revealed features unique to *M. polymorpha*. Methylation of CCG sites in *M. polymorpha* does not depend on *MET1*, unlike in *A. thaliana* and *Physcomitrella patens*. Furthermore, unlike *A. thaliana*, *M. polymorpha* shows higher methylation level at CAG sites than at other CHG contexts and CAG/CTG sites are mostly methylated asymmetrically. Interestingly, CAG and CTG methylation reached comparable levels and symmetry upon loss of CG methylation. Our results highlight the diversity of non-CG methylation regulatory mechanisms in plants.

## Introduction

Methylation of cytosine residues is a heritable epigenetic modification present in many eukaryotes, which influences transcription of genes and transposable elements (TEs). Most repetitive elements and TEs are highly methylated and transcriptionally repressed. TEs and their remnants residing near protein-coding genes (PCGs) can affect their epigenetic status and thus their expression. In many species, DNA methylation is also found in gene bodies. Although the function of gene body methylation is still controversial, it seems to positively correlate with gene transcription (Bewick and Schmitz 2017; Zilberman 2017).

In mammals, DNA methylation occurs predominantly at CG dinucleotides, whereas in plants, cytosine methylation is additionally present at non-CG contexts, including CHG and CHH motifs (where H is any base but G). Based on genetic approaches, multiple DNA methyltransferases were identified in *Arabidopsis thaliana*. METHYLTRANSFERASE 1 (MET1), an orthologue of the mammalian DNA METHYLTRANSFERASE 1 (Dnmt1), acts during DNA replication to maintain CG methylation. CHG methylation is mainly maintained by CHROMOMETHYLASE 3 (CMT3), while CHH maintenance requires the activity of CHROMOMETHYLASE 2 (CMT2) and the *de novo* DNA methyltransferase, DOMAINS REARRANGED METHYLASE 2 (DRM2). DRM2 is involved in the RNA-directed DNA methylation (RdDM) pathway, in which relies on 24-nt small interfering RNAs (siRNAs) and induces DNA methylation in all cytosine contexts.

Proteins of the MET1 family are well conserved within the plant kingdom. *A. thaliana met1* mutants with hypomethylation at CG sites display pleiotropic phenotype including delay in flowering, abnormal embryogenesis and seed abortion (Kankel et al. 2003; Saze et al. 2003; Xiao et al. 2006). In *Physcomitrella patens*, mutants of the *MET1* homologous gene *PpMET* showed defects in sporophyte formation (Yaari et al. 2015). Both *A. thaliana* and *P. patens met1* mutants display a drastic loss of methylation of cytosines in CG contexts, but also of the external cytosine of CCG trinucleotides, which represents one of the CHG subcontexts (Yaari et al. 2015; Zabet et al. 2017). Methylation at CCG depends on both MET1 and CMT3 orthologues, whereas methylation of other CHG subcontexts (CAG and CTG) only requires CMT3 (Yaari et al. 2015; Gouil and Baulcombe 2016). Thus, DNA methylases seem to cooperate, but precise analysis of DNA methyltransferases is still restricted to a few species and their relationships during plant evolution remain poorly understood.

In this report, we focus on one of the early land plants, the liverwort *Marchantia polymorpha*, that is in a basal lineage of land plants. *M. polymorpha* has peculiar characteristics with respect to DNA methylation. In particular, the DNA methylation status varies during *M. polymorpha* life cycle (Schmid et al. 2018). The observation of increased DNA methylation at all cytosine contexts in reproductive tissues (archegonia and antherozoids) and in the sporophyte implies epigenetic reprogramming in the germ cell lineage of *M. polymorpha*, unlike in other land plants. Furthermore, *M. polymorpha* lacks gene-body methylation in the gametophyte, before formation of the reproductive organs (Takuno et al. 2016). The absence of gene body methylation is also observed in other non-vascular plants including *P. patens*. Bewick et al. suggested that gene body methylation is restricted to angiosperms and that its establisment depends on CMT3 activity (Bewick et al. 2017). Athough non-vascular plants have CMT-like proteins, they are classified in a different clade than CMT2 and CMT3 and it has been proposed that this the lack of a functional ortholog of CMT3 can explain the absence of gene body methylation in non-vascular plants.

*M. polymorpha* contains a single copy of the *MET1* orthologous gene, Mp*MET*. Genetic nomenclature is as outlined in (Bowman et al. 2016). Mp*MET* is ubiquitously expressed throughout the development (Bowman et al. 2017; Schmid et al. 2018). We generated Mp*met* mutants and characterized their morphological phenotype and precise DNA methylation status. Disrupting Mp*MET* allowed us to dissect the controling mechanism of DNA methylation, including the relationships between CG, CHG, and CHH methylations and DNA methylation related proteins.

## Results

### Isolation of Mp*met* mutants

The domain structure of proteins of MET1 DNA methyltransferase family is highly conserved among plants (Pavlopoulou and Kossida 2007). N-terminal regions of MET1 proteins of *A. thaliana*, *P. patens*, and *M. polymorpha*, have two cytosine-specific DNA methyltransferase replication foci domains (DNMT1-RFD) (Fig S1), which are required for targeting the enzyme to the replication foci during DNA replication. MET1 proteins also have two bromo-adjacent homology (BAH) domains, which are assumed to act as protein-protein interaction modules and are commonly found in proteins that act in transcriptional silencing and chromatin remodeling (Kar et al. 2012). The DNA methyltransferase domain, located in C-terminus, has an enzymatic activity that is S-adenosylmethionine-dependent methyltransferases (SAM or AdoMet-MTase). AdoMet-MTases are enzymes that use S-adenosyl-L-methionine as a methyl group donor for the transmethylation reaction (Zhang and Zheng 2016). To analyze the function of Mp*MET* in *M. polymorpha*, we set about to generate mutants of Mp*MET* using the CRISPR/Cas9 system (Sugano et al. 2018; Sugano et al. 2014). We identified multiple mutants corresponding to three target regions (Fig. 1A, Fig. S2). Using the method of sporeling transformation (Ishizaki et al. 2008), two insertion and two deletion mutants were obtained, all containing a premature stop codon after the mutation site (Fig. S2). The Mp*met*-*1* and Mp*met*-*2* mutations were found in the second and third exon and located in first and second DNMT1-RFDs, respectively. Both Mp*met*-*3* and Mp*met*-*4* mutations were found in the 9th exon, and caused the deletion of a part of the DNA methyltransferase domain (Fig. 1A). All these Mp*met* mutant thalli (haploid gametophytic vegetative tissue) exhibited abnormal shape, reduced size and growth delay (Fig. 1B-D, Fig. S3). *M. polymorpha* forms an organ on dorsal side of the thallus for asexual reproduction, called the gemmae cup. Inside the gemmae cup, vegetative clones called gemmae are formed. However, Mp*met* mutants never formed neither gemmae cups nor gemmae. Furthermore, even under inductive conditions, neither female nor male Mp*met* mutants did not develop organs for sexual reproduction (archegoniophore and antheridiophore). After incubation on B5 media for more than a month, Mp*met* mutants generated many adventitious shoot-like structures and rhizoids on thallus (Fig. 1E). Dorsal structures, including air chambers and air pores, were formed but disorganized, and many undifferentiated cells were found on the surface of Mp*met* mutant thalli (Fig. 1F and G). Thalli of Mp*met* failed to develop a normal expansion pattern and lacked lateral growth. These observations revealed that Mp*MET* is necessary for promoting cell differentiation, maintaining differentiated cellular identities, or both in *M. polymorpha*.

**Fig. 1.**
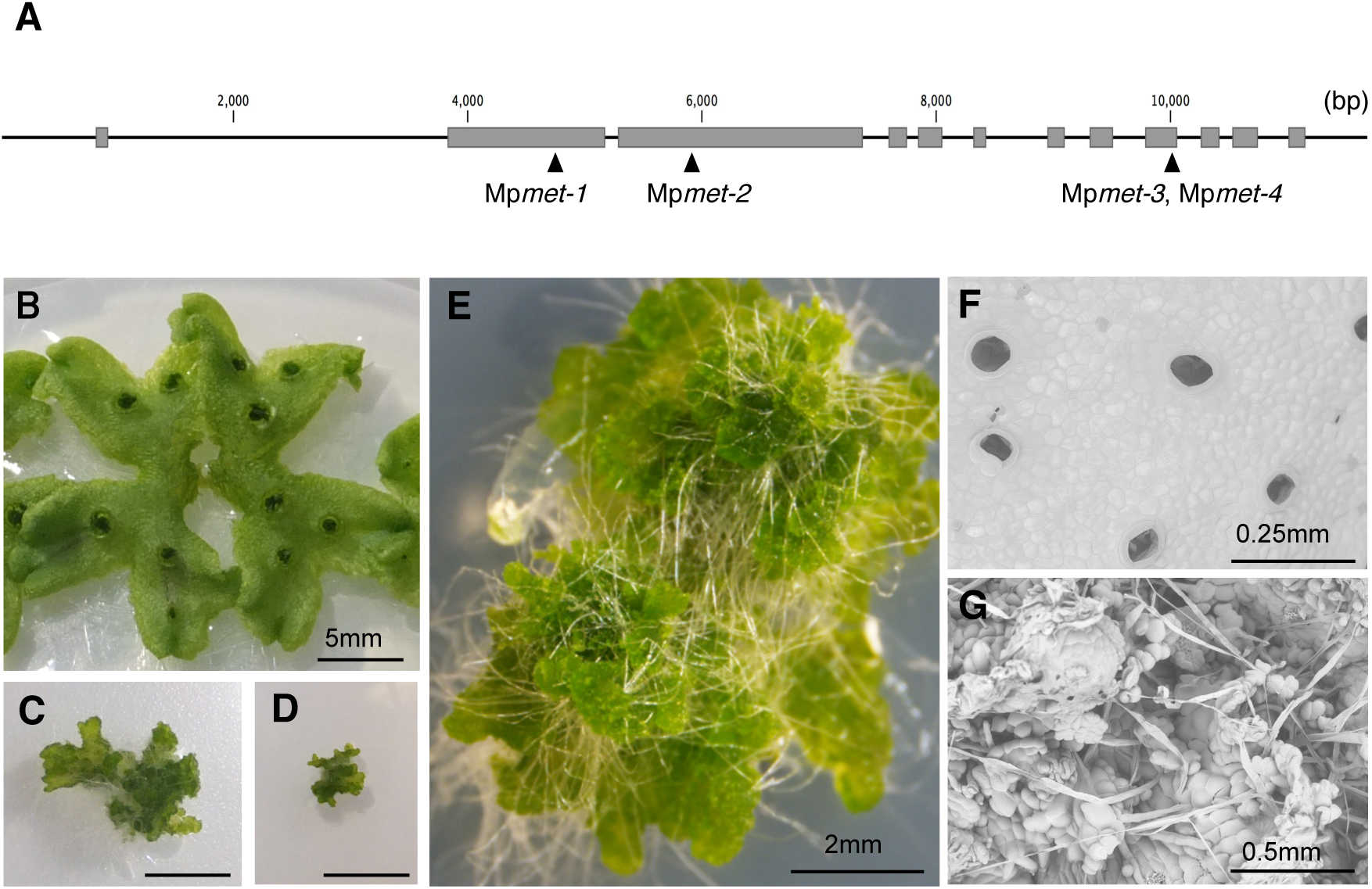
Developmental phenotypes of Mp*met* mutations in *M. polymorpha*. (A) Diagram showing the Mp*MET* gene in *M. polymorpha*. Mutation sites are represented by arrowheads. (B) 20-day-old WT Tak-1 grown on medium. Error bars; B-D 5mm, E 2mm, F 0.25mm and G 0.5mm. (C) 20-day-old Mp*met*-*1* mutant grown on medium. (D) 20-day-old Mp*met*-*2* mutant grown on medium. (E) 6-week-old Mp*met*-*3* mutant grown on medium. (F) SEM image of mature thallus of WT Tak-1. (G) SEM image of mature thallus of 6-week-old Mp*met*-*3* mutant.

We confirmed DNA hypomethylation of selected transposon-related regions in Mp*met* mutants using DNA methylation-sensitive restriction enzyme (Fig. S4). We analyzed DNA methylation levels of one specific copy of hAT-Ac family (Class II DNA transposon) and Ty3-gypsy (Class I retrotransposon), and the reduction of DNA methylation was observed in all Mp*met1* alleles.

### Multiple TEs are up-regulated in Mp*met* mutants

To identify genes and TEs regulated by Mp*MET*, we compared the transcriptomes of Tak-1 (wild type) and Mp*met* mutants generated by RNA-sequencing (RNA-seq). We used Tak-1 (male), Mp*met*-*3* (male) and Mp*met*-*4* (female) vegetative gametophytes (thalli). RNA-seq analysis was conducted in triplicates for the Tak-1 vs. Mp*met*-*3* comparison (Table S1) and in duplicates for the Tak-1 vs. Mp*met*-*4* comparison (Table S2). Among the 19,287 annotated genes, 1,447 and 225 were up-regulated in Mp*met*-*3* and Mp*met*-4, respectively. Of these, 1,260 and 38 genes were up-regulated specifically in Mp*met*-*3* and Mp*met*-*4* mutants, respectively. One hundred and eighty seven genes were commonly up-regulated in Mp*met*-*3* and Mp*met*-*4*, of which 92 genes were commonly expressed in Mp*met*-*3* and Mp*met*-*4* but not expressed in Tak-1 (Fig. 2A and Table S3a). The number of down-regulated genes in Mp*met*-*3* and Mp*met*-*4* was similar to that of up-regulated genes, but the number of genes that expressed only in Tak-1 (8 genes) was much less than those that were expressed only in Mp*met* mutants (Fig. 2A and Table S3b). Next, we categorized the commonly up- or down-regulated genes in Mp*met*-*3* and Mp*met*-*4* using GO terms (Alexa et al. 2006). Top 10 categories were picked up in Table S4, and various cellular events were included. Among up-regulated genes, terms of protein dimerization, methyltransferase activity, and metal ion and protein binding were enriched. These factors may be directly regulated by DNA methylation.

**Fig. 2.**
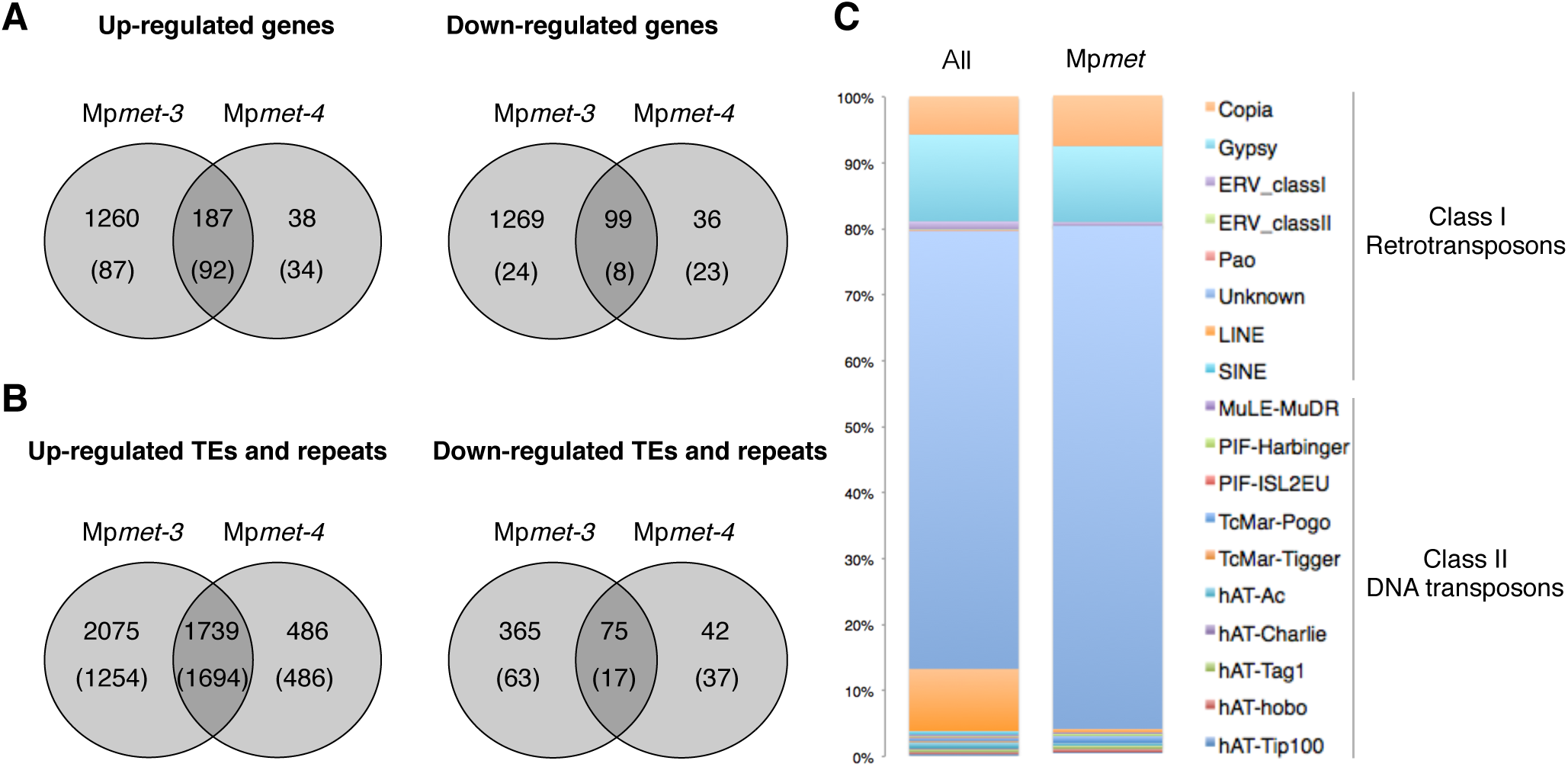
Summary of RNA-seq analysis of Tak-1, Mp*met*-3, and Mp*met*-*4*. (A) Venn diagram showing the number of differentially expressed genes in Mp*met*-3, and Mp*met*-*4* compare to Tak-1. The genes were selected using both >2 or <1/2 fold-change and a FDR of 0.05 after Benjamini-Hochberg correction for multiple-testing. The numbers in parenthesis show the number of genes for which reads were only detected in either Tak-1 or Mp*met*. In which case, the column of log2 (fold-change) in Table S1-S3 is represented as infinity (inf or -inf). (B) Venn diagram showing the number of differentially expressed TEs and repeats in Mp*met-3*, and Mp*met*-*4* compared to Tak-1. The differentially expressed TEs and repeats were selected using both >2 or <1/2 fold-change a FDR of 0.05 after Benjamini-Hochberg correction for multiple-testing. The numbers in parenthesis show the number of TEs and repeats for which reads were only detected from Tak-1 or Mp*met*. (C) Classification of TE-superfamily that are up- or down-regulated (total number is 1,814) in Mp*met*. The distribution of TEs in the *M. polymorpha* genome (All) is shown for comparison.

A total of 149,748 TEs and repeats have been annotated in the *M. polymorpha* genome (Bowman et al. 2017). We detected 1,739 TEs and repeats as being up-regulated in both Mp*met*-*3* and Mp*met*-*4;* most of these (1,694 TEs and repeats) were not detectably transcribed in Tak-1 (Fig. 2B and Table S5). On the contrary, only a much fewer number of TEs and repeats (75 TEs and repeats) were down-regulated in Mp*met*-*3* and Mp*met*-*4*. This indicates that CG methylation is important for silencing TE expression in *M. polymorpha*. Next, we categorized differentially expressed TEs into their respective superfamilies. More than 90% of *M. polymorpha* TEs consisted of class I retrotransposons, many of these are classified as unknown types (Fig. 2C). The proportion of retrotransposons and DNA transposons activated in Mp*met* mutants, was not different from that of the entire genome. However, the proportion of long interspersed nuclear elements (LINE) was very low in Mp*met* mutants compared to that of the entire genome.

### Genome-wide DNA methylation status in Mp*met* mutant

To reveal the function of Mp*MET* in DNA methylation, we performed genome-wide DNA methylation analysis using bisulfite-sequencing (BS-seq). Two genomic DNA samples extracted from Mp*met*-*3* vegetative gametophytes (Mp*met*_rep1, Mp*met*_rep2) and Tak-1 vegetative gametophytes (Tak-1_rep2) were treated with sodium bisulfite and analyzed by deep-sequencing (Table 1). We also included in the analysis the previously published BS-seq data of Tak-1_gametophyte (Bowman et al. 2017). In Mp*met* mutant, genome-wide CG methylation rate was decreased to around 0.5%, compared to Tak-1 (around 2.7%). On the other hand, non-CG methylation was dramatically increased in Mp*met* compared to Tak-1 at both CHG and CHH sequence contexts (Table 1). The results being very consistent between replicates, we selected the libraries with the highest sequencing coverage, Tak-1_gametophyte (hereafter referred to as WT) and Mp*met*_rep1 (hereafter called Mp*met*), for subsequent analyses.

**Table 1.**
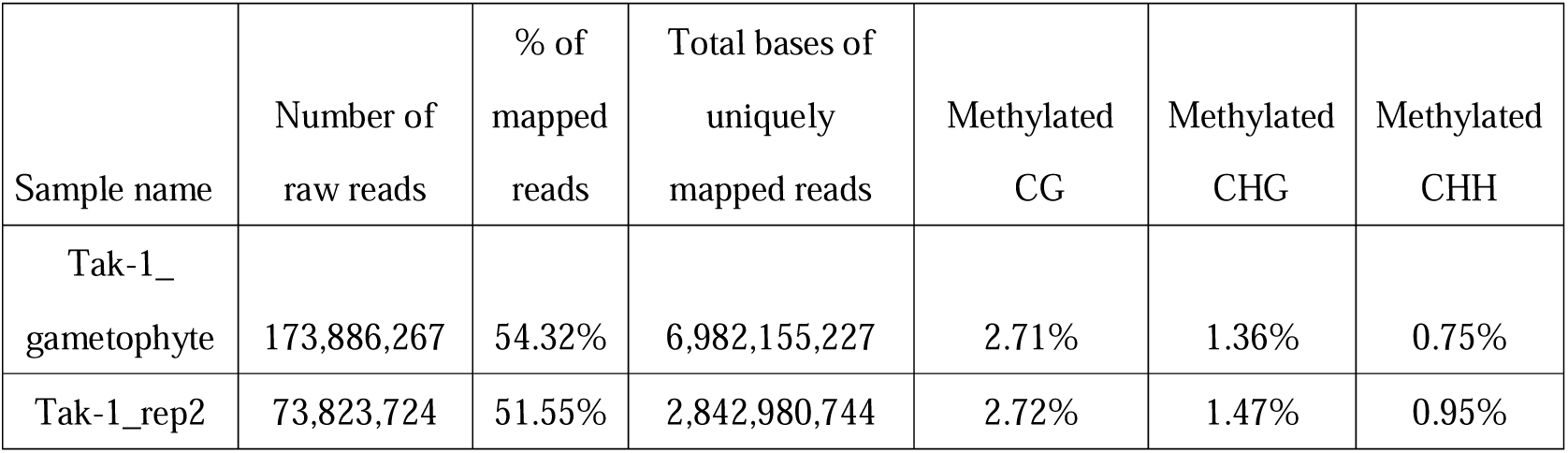
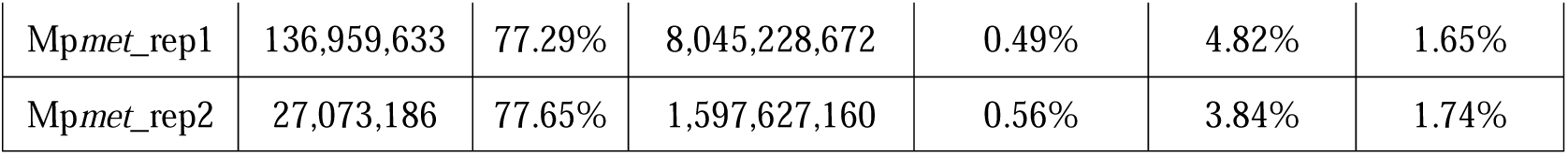
Summary of BS-seq analysis in this report.

Metaplot analyses of average DNA methylation over PCGs and TEs indicated that both types of loci lost CG methylation in Mp*met1* and that gene body CG methylation is very low in WT *M. polymorpha*, as previously reported (Fig. 3A, B) (Takuno et al. 2016). The average level of CG methylation over *M. polymorpha* TEs is lower than that of *A. thaliana* and cassava (35% vs. 80% and 90%, respectively) (Wang et al. 2015; Regal et al. 2016). Loss of *MET1* function in *A. thaliana* leads to contrasted effects on non-CG methylation at PCGs and TEs: while both PCGs and TEs show decreased CHH methylation in *met1*, TEs show global CHG hypomethylation whereas PCGs show increased CHG methylation in their bodies (Rigal et al. 2016). In *M. polymorpha* Mp*met* mutants, average levels of CHG and CHH methylation were strongly increased at TEs but not at PCGs. These results indicate that the pathways controlling DNA methylation differ between *A. thaliana* and *M. polymorpha*, and show a strong non-CG hypermethylation at TEs following depletion of CG methylation in *M. polymorpha*. Interestingly, CHG hypermethylation was less pronounced towards the central part of the TEs, whereas the increase in CHH methylation appeared rather homogenous along TEs. This suggests that hypermethylation at CHG and CHH sites in Mp*met* may occur through distinct regulatory pathways.

**Fig. 3.**
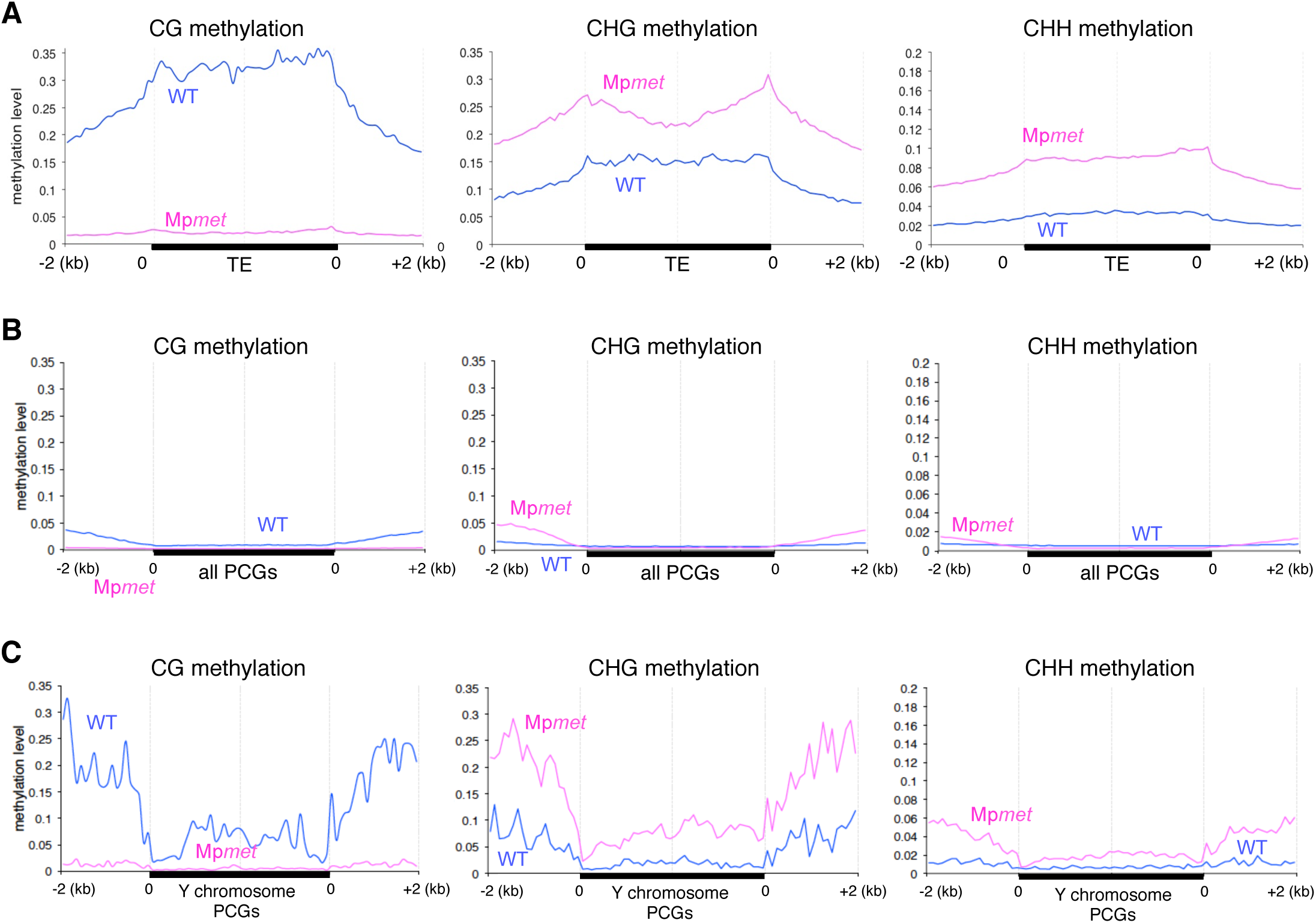
Average CG, CHG and CHH methylation over TEs (A), protein coding genes (PCGs, B) and PCGs located on chromosome Y (C) in WT Tak-1 and Mp*met* mutants. TEs (≥ 1 kb, n= 14,706), all PCGs (n = 22,318) and chromosome Y PCGs (n = 139) annotations were each divided in 40 bins of equal size. Their 5’ and 3’ 2 kb flanking regions were alignment and divided into 20 bins of 100 bp. Average methylation levels were computed for each bin and plotted.

In agreement with a previous report (Bowman et al. 2017), we found that PCGs located on the Y sex chromosome, unlike PCGs located on autosomes, showed a certain level of gene body CG methylation in the WT (Fig. 3C). In the Mp*met* background, CG methylation was virtually lost, while average levels of CHG and CHH methylation were increased over these genes, at both their bodies and flanking regions (± 2kb). Thus, changes of DNA methylation patterns of PCGs on the Y chromosome closely resemble those of TEs. This behavior is likely explained by the fact that PCGs on the Y chromosome are surrounded by, and in some cases contain repetitive elements and TEs in their bodies (Fig. S5).

To further investigate DNA methylation changes in Mp*met*, we compared the WT and Mp*met* methylomes and determined differentially methylated regions (DMRs) at CG, CHG and CHH cytosine sequence contexts (Fig. 4A). Confirming the Mp*MET* functions in CG methylation, the most abundant type of DMRs was hypomethylated at CG sites (CG hypo-DMRs), and we could not identify hypermethylated DMRs (hyper-DMRs) at CG sites. In comparison with hyper-DMRs, CHG and CHH hypo-DMRs were less numerous (2,382 and 5,019) and covered a much smaller fraction of the genome (2.6 and 5.6 Mb). DMRs showing CHG and CHH hypermethylation in Mp*met* were highly abundant (17,626 and 13,157), overlapping to a large extent with TE annotations and covering a larger fraction of the genome (22.2 and 21.3 Mb) similar to that of CG hypo-DMRs (20.2 Mb). Heat-map analysis showed that hypermethylation at CHG and CHH sites are well correlated in Mp*met* (Fig. 4B). Regions of non-CG hypermethylation in Mp*met* were not necessarily methylated at CG sites in the WT, and CG hypo-DMRs did not all gain methylation at CHG and CHH positions in Mp*met*. These results are consistent with the above-described data that non-CG methylation is highly increased at TEs in Mp*met1* and indicate that the presence of CG methylation in the WT is not a prerequisite for non-CG hypermethylation in Mp*met*.

**Fig. 4.**
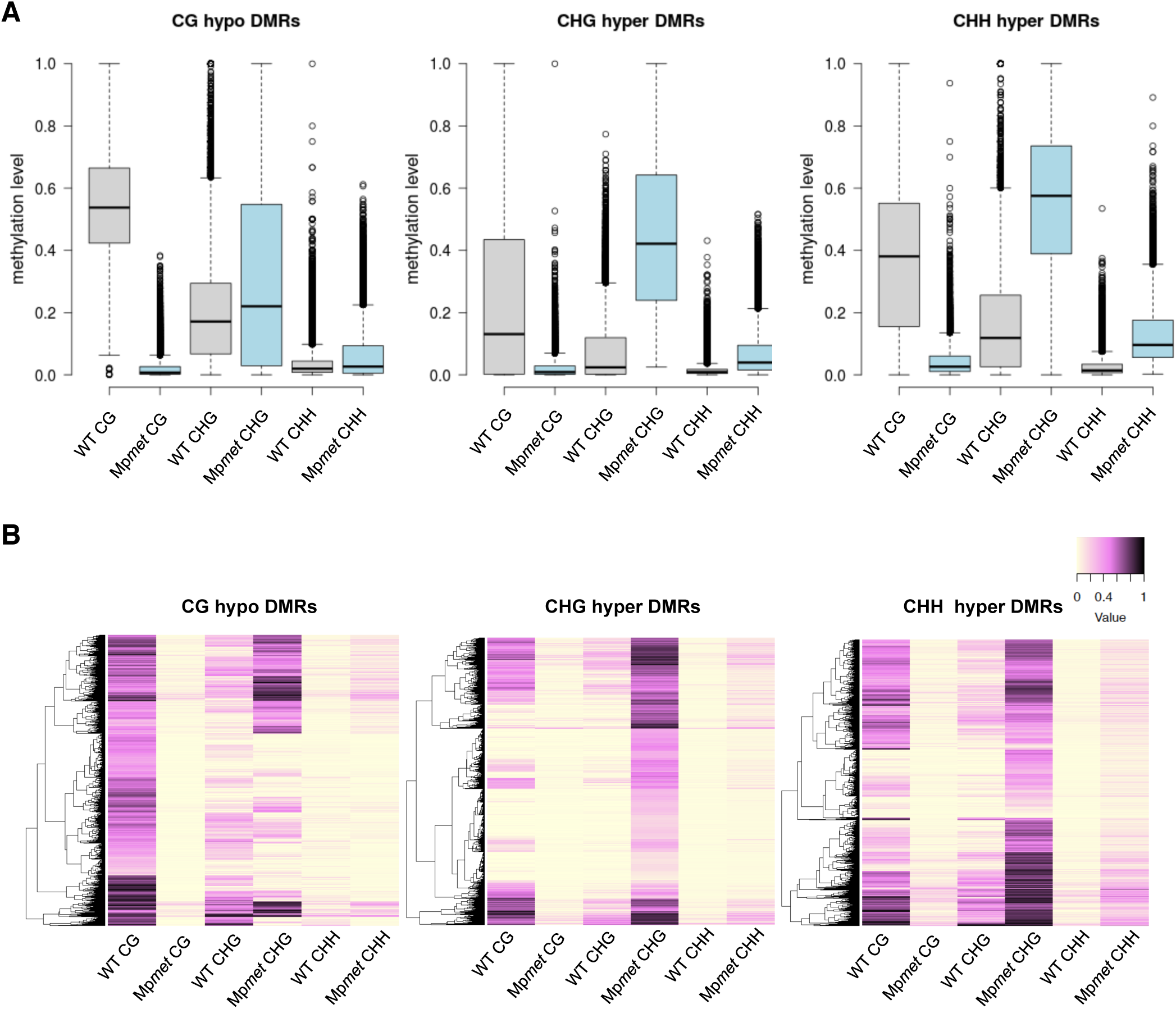
Differentially methylated regions (DMRs) in Mp*met* mutant. (A) Box plot of CG, CHG, and CHH methylation levels in WT and Mp*met* in CG hypomethylated regions (CG hypo DMRs), CHG hypermethylated regions (CHG hyper DMRs) and CHH hyper methylated regions (CHH hyper DMRs). (B) Heatmaps of CG, CHG, and CHH methylation levels in WT and Mp*met* at Mp*met* CG hypo DMRs, CHG hyper DMRs and CHH hyper DMRs.

### DNA methylation levels of non-CG trinucleotide subcontexts

Recently, it was reported that, in several plant species, DNA methylation levels differ depending on the trinucleotide sequence at CHG and CHH sites (Gouil and Baulcombe 2016). In *A. thaliana*, tomato, maize and rice, CAG/CTG sites appear more methylated than the external cytosine of CCG trinucleotides, at least in heterochromatin. In addition, it was shown in *A. thaliana* and *P. patens* that methylation of the external cytosine of CCG sites requires *MET1* in addition to *CMT3*, and is thus strongly reduced in corresponding *met1* mutants. Methylation levels also vary at CHH trinucleotide subcontexts, and one or two over-represented methylated CHH motifs are present in the plant species that were analyzed. For example, *A. thaliana* heterochromatin and the maize genomes are enriched with methylation at CAA and CTA sites, while CHH methylation in tomato heterochromatin is highest at CAA and CAT motifs. These differences in methylation at non-CG subcontexts in heterochromatin appear to result from a more efficient recruitment of CMT3 at CAG/CTG sites compared to CCG motifs, and a preference of CMT2 for CAA/CTA CHH sites (Gouil and Baulcombe 2016). Meanwhile, RNA-directed DNA methylation machinery contributes to non-CG methylation without any subcontext preference.

To get more insight into the mechanisms controlling DNA methylation patterns in *M. polymorpha*, we determined DNA methylation levels at non-CG trinucleotide subcontexts in the WT and in Mp*met*. Similar to other plant species analyzed so far, CHG methylation in WT *M. polymorpha* gametophytes was lower at CCG sites relative to CAG and CTG (Fig. 5A). However, while CAG and CTG methylation levels are indistinguishable in other plant species, CAG methylation level appeared higher than that of CTG methylation in *M. polymorpha*. This suggests that methylation is slightly more efficient at CAG sites than at CTG sites, and that CAG/CTG motifs are not methylated symmetrically (methylated on the two DNA strands at the same level). Indeed, examining methylation on the two DNA strands at CAT/CTG sites revealed that these sites largely tend to be methylated asymmetrically (only on one strand) in *M. polymorpha*, unlike in *A. thaliana* (Fig. 5B, Fig. S6, S7). This indicates that CAG/CTG methylation involves different molecular mechanisms in *A. thaliana* and *M. polymorpha*.

**Fig. 5.**
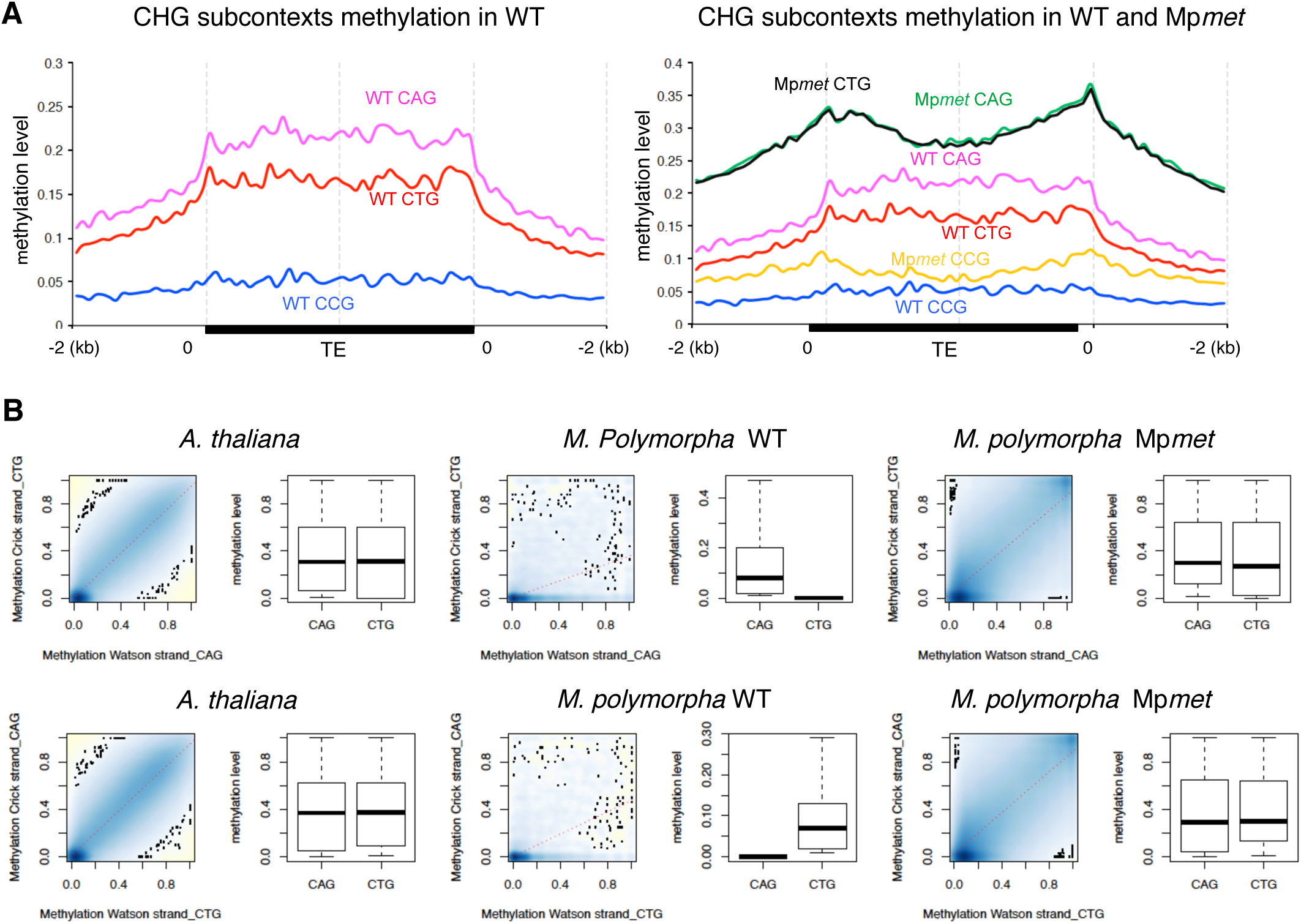
CHG methylation in WT and in the Mp*met* mutant. (A) Average methylation levels of CHG subcontexts (CAG, CTG and CCG) were plotted along TEs (≥ 1 kb) and their 2 kb flanking sequences as in Fig 3. (B) Scatterplots of methylation at CAGs (top) or CTGs (bottom) on the Watson DNA strand and of methylation at corresponding CA/TGs on the other DNA strand in *A. thaliana*, *M. polymorpha* WT and Mp*met* mutants. Only sites where cytosines were covered by at least 8 reads on each strand were retained in the analysis. Boxplots showing DNA methylation levels on both strands and are shown on the right of the corresponding scatterplots.

In Mp*met*, DNA methylation was increased at all three CHG subcontexts (Fig. 5A). This is in sharp contrast with the situation in *A. thaliana* and *P. patens*, where mutations in *MET1* lead to a strong decrease in CCG methylation (Gouil and Baulcombe 2016; Yaari et al. 2015). Remarkably, CAG and CTG sites reaches similar levels of methylation in Mp*met* and CAG/CTG methylation became symmetrical. These results reinforce the notion that pathways maintaining CHG methylation patterns in *M. polymorpha* and *A. thaliana* are different. These results further suggest that, in the Mp*met* background, CAT/CTG methylation might be taken over by a different pathway than in WT gametophytes.

The DNA methylation of CHH subcontexts in *M. polymorpha* differs from other species analyzed to date (Fig. 6). In heterochromatin of *A. thaliana* and maize, CHH methylation levels are highest at CAA and CTA sites; CAA and CAT sites are preferentially methylated in tomato, while rice shows more methylation at CTA positions (Gouil and Baulcombe 2016). Analyzing CHH methylation along TEs in wild-type *M. polymorpha* revealed a preference for methylation at CHH motifs with a cytosine in the second position (CCC, CCT and CCA). Since the CHH methylation biases reported in other plant species are influenced by various chromomethyltransferases, the differential methylation at CHH subcontexts may be mediated by one (or both) of the chromomethylase encoded by the *M. polymorpha* genome. In Mp*met* CHH methylation increased at all nine CHH subcontexts, although to various extend. As a result, CHH methylation preference was changed in Mp*met* and TEs showed highest methylation at CTA sites. This suggests that both RdDM and a CMT-mediated CHH methylation pathways are altered in Mp*met*.

**Fig. 6.**
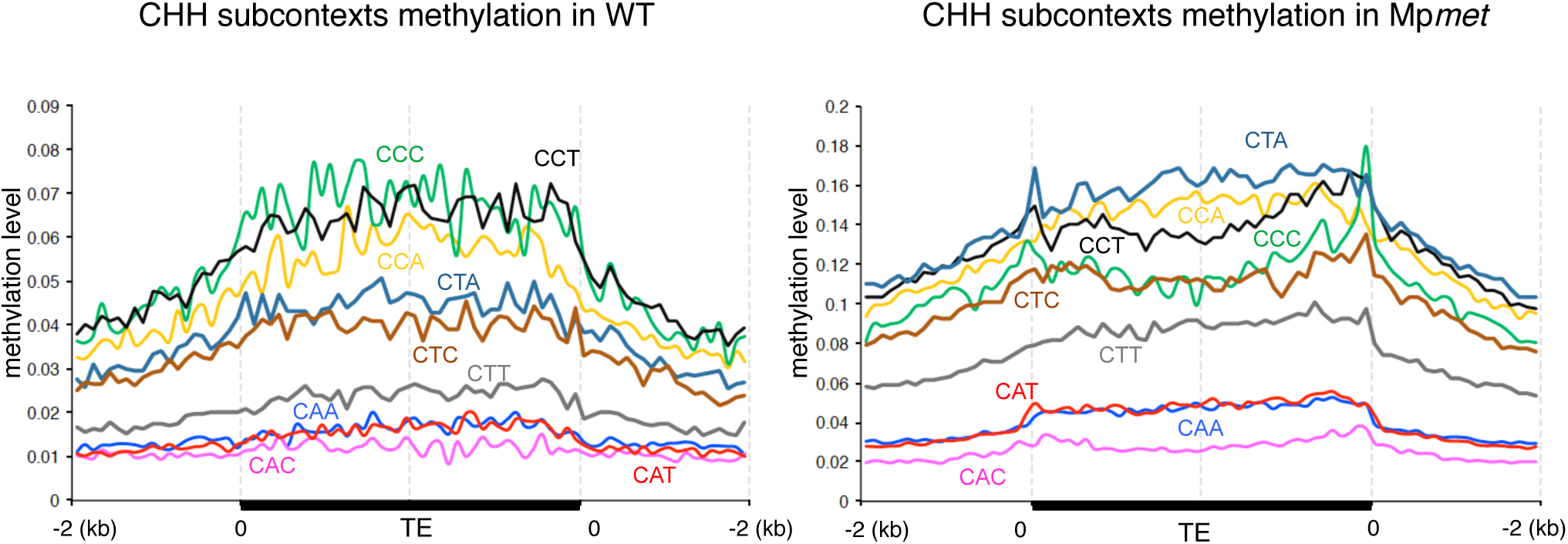
CHH methylation in WT and in the Mp*met* mutant. Average methylation levels of CHH subcontexts were computed and plotted along TEs (≥ 1 kb) and their 2 kb flanking sequences as in Fig 3.

## Discussion

### Mp*MET* is required for maintaining differentiated cellular identities in vegetative gametophyte and formation of the sexual reproductive organs

Here, we generated multiple Mp*met* mutant alleles, which showed severe morphological defects. The vegetative gametophytes of Mp*met* mutants lost tissue differentiation except for rhizoid production. Our observations suggest that Mp*MET* plays an important role in maintaining differentiated cellular identities during multicellular gametophytic growth. Although the *M. polymorpha* Mp*met* mutants have common morphogenic phenotypes during vegetative stages, mutants of the *P. patens MET1* orthologous gene do not show any strong defects during gametophyte differentiation (Yaari et al. 2015). In *A. thaliana*, several genes are known to be regulated by CG methylation in specific biological phenomenon. Hence, particular genes controlling cell differentiation may be regulated by Mp*MET*. It is known that auxin promotes cell differentiation in *M. polymorpha*, and auxin-insensitive plants, or mutants defective for auxin biosynthesis genes, exhibit a phenotype similar to Mp*met* mutant (Flores-Sandoval et al. 2015; Kato et al. 2015). Therefore we looked in Mp*met* for possible expression changes of genes known to be involved in auxin biosynthesis and signaling pathway (Table S6) (Eklund et al. 2015; Flores-Sandoval et al. 2015; Kato et al. 2015). We found down-regulation of the auxin biosynthesis gene *MpYUC2*, and major auxin response genes tend to be slightly down-regulated in Mp*met* mutants. However, the expression change is rather moderate, and the link between the phenotype of Mp*met* vegetative gametophyte and changes in the expression of auxin related genes needs further investigation.

In *P. patens*, *Ppmet* mutant shows defects only during sporophyte development (Yaari et al. 2015), whereas the *M. polymorpha Mp*met** mutants never developed neither archegoniophore nor antheridiophore. Mp*MET* is expressed throughout the life cycle, with a relatively higher expression in archegoniophore and antheridiophore (Bowman et al. 2017; Schmid et al. 2018). These findings indicate that Mp*MET* has an important role for in the transition between the vegetative and reproductive stages..It was reported that DNA methylation patterns are dramatically different between sporophytes, vegetative gametophyte, and gametangia in *M. polymorpha*, suggesting the existence of epigenetic reprogramming at least twice in the life cycle (Schmid et al. 2018). Mp*MET* may have functions in many specific stages of the life cycle. In this report, we used Mp*met* mutants and focused on the function in gametophyte, but conditional knock out and reporter analysis may elucidate more precise function of Mp*MET* throughout the life cycle.

### Non-CG methylation is strongly facilitated in Mp*met* mutant

Our methylome analysis revealed that Mp*met* mutants display a unique DNA methylation pattern in that non-CG methylation levels are drastically increased at TEs. Such increase in non-CG methylation has not been observed in *Ppmet* mutants of *P. patens* (Yaari et al. 2015). In *A. thaliana*, while PCGs tend to gain CHG methylation in their body in first-generation *met1* mutants, TEs do not show increased non-CG methylation, but rather decreased average CHG and CHH methylation levels (Rigal et al. 2016). Aberrant *de novo* non-CG methylation was detected in late generation selfed *A. thaliana met1* mutants (Mathieu et al. 2007). The alterations in non-CG methylation in *A. thaliana met1* mutants have been attributed to down-regulation of *ROS1* and *IBM1*, as well as to increased DRM2-mediated RdDM associated with increased accumulation of 24-nt siRNAs at some repeated loci (Mathieu et al. 2007; Rigal et al. 2016; Rigal et al. 2012). Comparing transcript accumulation of the *ROS1*, *IBM1* and *DRM2* orthologs in WT *M. polymorpha* and Mp*met* mutants revealed no significant variation, apart from a slight increase in MpDRMa expression (Table S6). Therefore, it is likely that the alterations in non-CG methylation in Mp*met* and *A. thaliana met1* mutants occur through distinct molecular mechanisms. Making the analogy with other plant systems, we propose that at least part of the increased DNA methylation in Mp*met* involves a RdDM-like pathway since it affects both CHG and CHH sites. Hence, our data suggest that CG methylation (and/or Mp*MET*) strongly inhibit RdDM in *M. polymorpha*.

### *M. polymorpha* shows a specific feature of DNA methylation of CHG and CHH subcontext

In *A. thaliana*, tomato, maize and rice, CCG methylation is 20-50% lower than CAG/CTG methylation, at least in heterochromatin, and we show here that it is also the case in *M. polymorpha*. However, our results indicate the regulation of CHG methylation in *M. polymorpha* likely involves distinct/unique molecular mechanisms. Indeed, while methylation of the external cytosine at CCG sites requires *MET1* in *A. thaliana* and *P. patens* (Gouil and Baulcombe 2016; Yaari et al. 2015), we found that it is not the case in *M. polymorpha*, in which Mp*MET* rather seems to inhibit CCG methylation. In addition, we found that, unlike in other plant species analyzed so far, methylation levels at CAG and CTG sites are not equivalent in *M. polymopha*, and that methylation at these sites is mostly asymmetrical. In *A. thaliana*, it has been proposed the difference between CCG and CAG/CTG methylation levels is due to preferential recruitment of CMT3 to CAG/CTG sites by histone methylase Su(var)3-9 homologue (SUVH) family protein SUVH4, while recruitment of CMT3 to CCG is less efficient and depends on SUVH5/6 (Gouil and Baulcombe 2016). The *M. polymorpha* genome encodes only one SUVH-like protein and two CMT-like proteins that reside in an independent clade, outside of the CMT3 and CMT2 clade (Bewick et al. 2017; Bowman et al. 2017). It is possible that the *M. polymorpha* SUVH-like and/or CMT-like proteins are responsible for the differential methylation at CAG, CTG and CCG sites. Because CHG methylation is largely asymmetrical in *M. polymorpha*, we propose that it is mostly maintained by a DNA replication independent RdDM-like pathway, rather than by a CMT3-like activity. Although methylation was not analyzed for individual CHG subcontexts, it has been observed recently that CHG methylation is asymmetrical in the flowering plant *Eutrema salsugineum* (Bewick et al. 2016). Consistent with our hypothesis in *M. polymorpha*, *E. salsugineum* genome also seems to lack *CMT3*. The RdDM-like pathway regulating CHG methylation in *M. polymorpha* may involve the DRM-like but also the CMT-like proteins as effectors, which may be at the origin of the differential CHG subcontext methylation.

We report that CHH methylation of TEs shows a subcontext preference for CCC, CCT and CCA in WT *M. polymorpha*. This bias differs from the ones that have been observed in the plant species analyzed so far, and which have been assigned to originate from CMT2-like activities. This CHH subcontext methylation bias is also likely reflecting the action of the *M. polymorpha* CMT-like proteins. The fact that this CHH subcontext preference differs from other plant species analyzed so far is consistent with the fact that CMT-like proteins encoded by the *M. polymorpha* genome are in a separate clade than CMT2.

In the Mp*met1* mutant background, the increase in DNA methylation affects all non-CG contexts, suggesting that RdDM-like pathway is stimulated in the absence of CG methylation. Remarkably, the hypermethylation is not merely an increase in pre-existing WT non-CG methylation levels, and patterns of non-CG methylation are modified: CAG and CTG methylation reach similar levels and symmetry in Mp*met* mutant, and the CHH subcontext methylation preference differ from that of the WT. Symmetry of CHG methylation in Mp*met* suggests that it is maintained in a DNA replication-coupled manner in this mutant background, potentially by a CMT3-like activity, which would be repressed by CG methylation and/or Mp*MET* in the WT. In addition, the changes in CHH subcontext methylation preference also argue that different CMT-like activities act in WT and Mp*met* plants. Analyzing methylomes of all mutants of the various DNA methyltransferases and DNA methylation related proteins in *M. polymorpha* will certainly provide useful insights into the unique regulatory mechanisms of DNA methylation.

In summary, our analysis of WT and Mp*met* mutant methylomes reveals features unique to *M. polymorpha*, and highlight that regulatory mechanisms of non-CG methylation are not conserved during land plants evolution. Further analysis of the methylomes in mutant backgrounds for the various DNA methyltransferases and DNA methylation-related proteins of *M. polymorpha* is now needed to provide new insights into the unique regulatory mechanisms of DNA
methylation in this emerging model organism.

## Materials and Methods

### Plant materials and growth condition

*M. polymorpha* male accession Takaragaike-1 (Tak-1) was used as WT in this study. F1 spores generated by crossing between Tak-1 and female accession Takaragaike-2 (Tak-2) were used to obtain Mp*met*-*1* and Mp*met*-*2*, and the BC4 spores generated by crossing between Tak-1 and a female progeny of Tak2 backcrossed to Tak-1 for three times (BC3-38) were used to obtain Mp*met*-*3* and Mp*met*-*4 alleles* (Bowman et al. 2017; Yamaoka et al. 2018). *M. polymorpha* was grown on half-strength Gamborg’s B5 medium containing 1% sucrose and 1% agar at 22°C under continuous white light.

### Targeted mutagenesis by CRISPR/Cas9 in *M. polymorpha*

Mp*met* mutant alleles were obtained by CRISPR/Cas9 system optimized for *M. polymorpha* by Sugano et al.(Sugano et al. 2018) Oligo DNA corresponds to three single-guide RNA (sgRNA) target regions (5’-CTTGGTCCTGTCCACTCG-3’, 5’-AGTAT CTT GGTACCGGCT-3’ 5’-CGATCTTCCTGCCGAGA-3’, respectively) were inserted into pMpGE_En03 vector including U6 promoter within BsaI restriction site. Each entry vector was recombined with pMpGE010 destination vector. These constructed binary vectors were transformed to spores that were written in plant materials, using *Agrobacterium tumefaciens* GV 3101 strain with the method by Ishizaki et al (2008).

### Scanning electron microscopy

Microscopy images were taken with Hitachi Miniscope TM3000 (Hitachi High-Technologies Corp., Tokyo, Japan).

### DNA methylation assay using DNA methylation-sensitive restriction enzymes

Genomic DNA was extracted from *M. polymorpha* thallus by Wizard Genomic DNA purification kit (Promega) according to the manufacturer’s instructions. 100 ng of genomic DNA was digested with 20U of DNA methylation sensitive enzymes, HpaII and McrBC (TaKaRa) for 12h at 37°C. After digestion, enzymes were inactivated by 70°C 20min. Digested and undigested DNAs (2ng) were then amplified by 35 cycles of PCR using following primers; hAT-Ac (7_hAT-Ac_F1, 5’-TTACTTTCCACCACCACATAATCAATGT-3’ and s7_hAT-Ac_R1, 5’-CGT GAAGAGGCATAGAT GGGTCGAGACT-3’), Ty3-gypsy; (S112_LTR 1-McrBC-F1, 5’-TTACGAGTCGGCTCACGAAGTTC-3’ and S112_LTR1 -McrBC-R1, 5’-AACACCTCTGATCGCTCCATCTAT-3’).

### RNA-seq analysis

Total RNA was extracted by RNeasy kit (QIAGEN) according to the manufacturer’s instructions. RNA was quantified by Invitrogen Qubit 3 fluorometer and Qubit HS RNA assay kit (Thermofischer scientific). Library preparation by TruSeq RNA sample preparation kits (Illumina) and sequencing by Hiseq1500 (Illumina) were descrived previously. The sequence reads were mapped to the *M. polymorpha* genome sequence v.3.1 by TopHat v.2.0.13 with default parameters. The mapped reads were used to identify the differentially expressed genes between WT and Mp*met* by calculating FPKM values using Cuffdiff v.2.1.1. Three biological replicates for WT vs Mp*met*-*3* and two biological replicates for WT vs Mp*met*-*4* were used for RNA-seq analysis. Differentially expressed genes and TEs were identified after Benjamini-Hochberg correction for multiple-testing within each replicates.

### BS-seq analysis

Genomic DNA was extracted using standard CTAB-procedure, and cleaned using a DNA purification kit (Zymo Research). One ug of genomic DNA was sheared using Covaris S220 Adaptive Focused Acoustics ultra sonicator. Libraries were constructed following standard protocols using the NEBNext Ultra II DNA library preparation kit (NEB) and Illumina-compatible adaptors in which all cytosines were methylated. The ligated adaptors were treated with sodium bisulfite using the ZYMO DNA methylation lightning kit (Zymo Research), and the libraries amplified with the Thermo Fisher Phusion U Hot Start DNA polymerase (Thermo Fisher Scientific). Samples were barcoded, and sequenced using a NextSeq500 machine at the CSHL genome Center. The sequence reads were first filtered using TRIMMOMATICS (http://www.usadellab.org/cms/?page=trimmomatic). Quality was verified using FASTQC, and the cleaned reads mapped to the M. polymorpha genome sequence v.3.1 using BISMARK with the following parameters: score_min L,0,-0.4.

### Identification of differentially methylated regions (DMRs)

DMRs between WT and Mp*met* were separately called for CG, CHG and CHH sequence contexts using a dynamic fragmentation strategy (Guo et al. 2018). Only cytosines covered by at least 4 reads in both samples were considered. Background fragments were dynamically defined as containing at least 5 cyotsines, a maximal distance of 400 bp between two adjacent cytosines and a maximal length of 2000 bp. Shared cytosines within each background fragment were deemed differentially methylated using an unpaired *t*-test (*P* ≤ 0.01) and we defined DMRs using a methylation rate difference of at least 40% for CGs, 20% for CHGs and 10% for CHHs.

### Funding information

Y. I. was supported by the Program to Disseminate Tenure Tracking System, MEXT, Japan, the Japan Society for the Promotion of Science [grant number 15K18578], and Wesco Scientific Promotion Foundation. D.G. was supported by a Marie Sklodowska-Curie individual fellowship (658900). M.A.A.V. was supported by Consejo Nacional de Ciencia y Tecnología CB158550, Cuerpo Académico CA-UVER-234, Agropolis Fondation EPIMAIZE, Royal Society Newton Advanced Fellowship NA150181, and a Royal Society Research Fellowship (RG79985). A.A.C. is the recipient of a graduate scholarship (629316) from CONACYT. T. K. was supported by NIBB Collaborative Research Programs number 16-425. Work in the O. M. laboratory was supported by CNRS, Université Clermont Auvergne and Inserm and ERC grant I2ST 260742.

### Disclosures

No conflicts of interest declared.

## Acknowledgments

We would like to thank S. Sugano for CRISPR/Cas9 vectors, S. Shigenobu for RNA-seq, Ms S. Ishida for *M. polymorpha* transformation, and M. Kato, M. Fujii, and M. Uemura for technical assistance.

**Supporting information/ additional files (if appropriate):**

**Supplementary data** – **6 supplementary Tables and 7 supplementary figures.**

